# A liquid chromatography-mass spectrometry method to quantify total Coenzyme A concentration and isotopic labeling

**DOI:** 10.64898/2026.05.19.726225

**Authors:** Amelia L. Taylor, Nathaniel W. Snyder, Caroline R. Bartman

## Abstract

Coenzyme A is an essential cofactor synthesized from pantothenate, cysteine, and ATP, and is involved in numerous processes of cellular metabolism through its ability to carry activated acyl groups. Coenzyme A participates in catabolism of carbohydrate, fat and amino acids; biosynthesis of fatty acids, cholesterol and heme; and protein modification including acetylation and 4-phosphopantetheinylation. Despite CoA’s critical functions, the regulation of CoA levels and the rate of CoA synthesis in different cell types and disease states are not well understood. One reason for this gap is that many acyl-CoA species are analytically challenging to measure due to factors including instability, poor ionization, and the wide range of biochemical properties conferred by different acyl chain lengths. In addition, most current methods do not support analysis of CoA isotopic labeling, which is required to quantify CoA synthesis rate or to measure absolute concentration using isotope-labeled internal standards. Here, we describe a method to quantify the concentration and isotopic labeling of total CoA, defined as the sum of CoASH plus all acyl-CoA species. Acyl-CoA species are hydrolyzed using sodium hydroxide to remove acyl chains, then CoA is derivatized on the thiol with N-ethylmaleimide (NEM). Following protein precipitation and solid phase extraction, samples are analyzed by liquid chromatography-mass spectrometry. This method is linear in a wide range that captures mouse tissue CoA levels, with accuracy within 15% error and precision below 15% relative standard deviation for both pure standards and tissue samples. We applied this method to measure total CoA concentration in five tissues from male and female mice, and total CoA synthesis rate in mouse liver via infusion of ^13^C-^15^N-pantothenate. Overall, this method offers a tractable approach to measure total CoA concentration and isotopic labeling to enable study of total CoA synthesis rates and concentrations in health and disease.

## Introduction

Coenzyme A is an essential cofactor synthesized from Vitamin B5 (also known as pantothenate), cysteine, and ATP and is involved in numerous processes of cellular metabolism.^1,2^ Coenzyme A is a carrier of activated acyl groups and participates in catabolism of carbohydrate, fat and amino acids; anabolism of fatty acids, cholesterol and heme; and protein modification including acetylation and 4-phosphopantetheinylation (Figure 1A).^3–6^ The concentration of Coenzyme A in cells can govern the ability to carry out these reactions. Mutations in the CoA synthesis pathway lead to neurodegenerative and cardiac disorders in human patients.^7^ In addition to these monogenic disorders, alterations in CoA metabolism are proposed to associate with cancers and cardiovascular disease.^8–10^ Due to Coenzyme A’s role in health and disease, there is strong interest to the field in quantitative methods to measure CoA concentration and synthesis rate.

**Figure 1.**
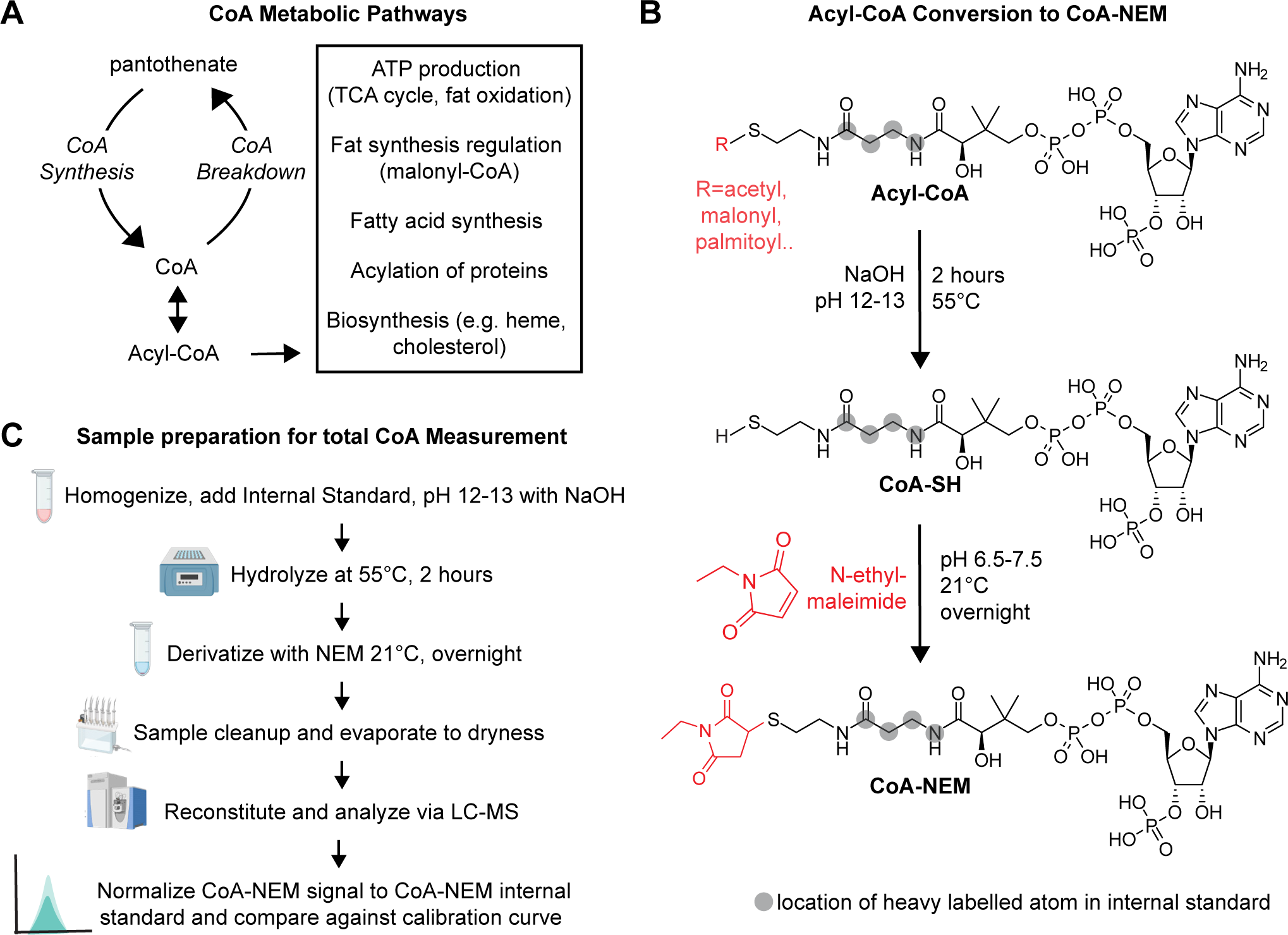
Sample preparation and derivatization strategy for total CoA measurement. **A)** Schematic of Coenzyme A metabolic functions. **B)** Hydrolysis and N-ethyl-maleimide (NEM) derivatization reaction for acyl-CoA to CoASH and then CoA-NEM. Gray circles indicate locations of stable isotope labels in internal standard retained through sample preparation. **C)** Sample preparation workflow for total CoA measurement by liquid chromatography-mass spectrometry including acyl-CoA hydrolysis and N-ethyl-maleimide derivatization.

CoA exists in the unconjugated form (known as CoASH) as well as various acyl-CoA forms, including a wide variety of acyl chain lengths from acetyl- to palmitoyl- and beyond. To measure cellular CoA concentration and synthesis rate, one could measure each individual species (CoASH and each acyl-CoA), or one could measure the total CoA backbone from all these species by hydrolyzing off the acyl chains and measuring CoASH or derivatized-CoASH. Measuring total CoA has several benefits. First, the different CoA species have very different biochemical properties due to different acyl chain lengths, so no single method is able to capture all of them without complicated separations.^11^ Second, many CoA species can be challenging to measure because of low concentration (e.g. CoASH) or propensity to spontaneously hydrolyze (succinyl-CoA). Third, since the CoA backbone of CoASH and acyl-CoAs are in fast exchange due to high fluxes of CoA-dependent pathways, capturing the synthesis rate of the CoA backbone is best carried out by measuring total CoA.

Current approaches to measure total CoA fall into the following categories: (A) enzymatic spectroscopy, (B) enzymatic radioassays, (C) high performance liquid chromatography (HPLC)-UV/Vis spectroscopy or fluorescence detection (FLD), or (D) liquid chromatography-mass spectrometry (LC-MS) ^12–24^. Most of these methods, while sensitive, lack the ability to capture the total CoA content of the cell, instead reporting a summed measurement of the most abundant short-chain CoA species (e.g. acetyl-CoA, malonyl-CoA, CoASH). Additionally, enzymatic and spectroscopic methods are unable to distinguish isotope-labeled CoA from native CoA, making the measurement of CoA synthesis using stable-isotope-labeled precursor metabolites impossible and requiring the use of external rather than internal standards for quantification.

In this paper, we set out to develop and validate an LC-MS method to measure total CoA concentration and isotopic labeling. In contrast to the enzymatic or fluorescent assays generally used to measure total CoA, a mass-spectrometry measurement method will ensure the molecular specificity of the measurement by measuring mass-to-charge ratio, and will enable measurement of heavy-isotope-labeled CoAs, either as internal standards or introduced from labeled precursors to quantify Coenzyme A synthesis. To isolate total CoA, we used heat under basic conditions to hydrolyze acyl chains, derivatized the resulting CoASH with N-ethyl-maleimide, then carried out solid phase extraction and LC-MS measurement. We tested the method’s limit of quantification, linearity, accuracy, precision, carryover, stability, and matrix effect and report figures of merit according to the guidelines recommended by the FDA’s bioanalytical method validation guidance for industry.^25^ Overall, this method enables LC-MS measurement of total CoA concentration and isotopic labeling with high precision and accuracy.

## Methods

### Standards and Chemicals

Acetyl-Coenzyme A lithium (HY-113596A) and succinyl-Coenzyme A sodium (HY-148285) were obtained from MedChemExpress. Palmitoyl-CoA (16:0-CoA) was obtained from Avanti Polar Lipids (870716P). N-ethyl-maleimide was obtained from Fisher Scientific (AA4052603H). ^13^C_3_-^15^N_1_ stable isotopically-labeled pantothenate was purchased from Cambridge Isotope Laboratories (CNLM-7694). Glacial acetic acid (Millipore Sigma, AX0073-9), Trizma-HCl (Millipore Sigma, T3253), DMSO (Fisher, BP231), and sodium hydroxide (NaOH, Fisher, SS277) were obtained from Millipore Sigma or Thermo Fisher Scientific. Optima LC/MS-grade water (Fisher, W5-4), methanol (MeOH, Fisher, A454-4), ethanol (EtOH, Fisher, AC611050040), ammonium hydroxide (Honeywell, 60-003-09), ammonium formate (Millipore Sigma, C951V20), ammonium phosphate (Fisher, 01-337-485), methylenediphosphonic acid (Millipore Sigma, C931D73), ammonium acetate (Sigma, 09689), and acetonitrile (ACN, VWR, NC1641805) were obtained from Thermo Fisher Scientific, VWR International or Millipore Sigma. Fetal bovine serum was obtained from Millipore Sigma (26400044, lot 23M233).

Standard isotopically-labeled acyl-Coenzyme A mix was prepared in-house, using the previously described protocol.^26^ Briefly, Pan6-deficient yeast cells were grown in media with ^13^C_3_-^15^N_1_ stable isotopically-labeled pantothenate (Cambridge Isotope Laboratories, #CNLM-7694) in place of unlabeled pantothenate for 31 hours, from which a standard isotopically-labeled acyl-CoA mix was extracted and purified.^26^ This standard was aliquoted and stored at - 80°C until use.

## Mouse Studies

All mouse studies were approved by the Institutional Animal Care and Use Committee of the University of Pennsylvania. Quantitation of tissue CoA was performed in n=5 male and n=5 female 11-week-old C57BL/6 mice from Charles River Laboratories. Mice were maintained under a 12-hour light-dark cycle (7am-7pm). On the day of tissue collection, mice were fasted at 7 AM yet left with continuous access to water. At 4 PM, mice were euthanized by cervical dislocation, and tissues were immediately harvested and freeze-clamped using a Wollenberger clamp cooled in liquid nitrogen. Tissues were stored at -80°C until measurement (see Tisue Extraction and Preparation below).

For measurement of CoA synthesis rates, LSL-Trp53^R172H/+^, Pdx-1-Cre mice were used. These mice are littermates to a pancreatic cancer genetically-engineered mouse model and have one mutant allele of p53 but are healthy and cancer-free.^27^ Aseptic surgical techniques were used to place a catheter (Instech, C20PU-MJV2011) in the right jugular vein and to connect the catheter to a vascular access button (VABM1B/25, Instech) implanted under the back skin of the mouse. Mice were allowed to recover for three to five days, during which they were monitored daily. ^13^C_3_-^15^N_1_ stable isotopically-labeled pantothenate (Cambridge Isotope Laboratories, CNLM-7694) at a concentration of 0.1mM in sterile saline was intravenously infused at a rate of 0.1µL/g body weight/min using World Precision Instruments AL300 rodent infusion pumps for 24 or 48 hours. During infusions, mice were housed in a private procedure room within the mouse facility with a 12-hour light/dark cycle matching the original housing room, with food and water provided *ad libitum*. Blood was collected via tail snip at 0, 24, and 48 hours and immediately placed on ice. Whole blood was centrifuged for 10 minutes at 21,300xG, and serum was aliquoted and stored at -80°C until further use. Tissues were harvested as above.

### Total CoA sample preparation

Calibration and quality control samples were prepared by adding purified standards (acetyl-CoA, MedChemExpress, HY-113596A) to 60% fetal bovine serum (Millipore Sigma #26400044, lot 23M233) as a pseudo-matrix in a final volume of 100 µL. For mouse tissue samples, clamp-frozen whole tissue samples were transferred to 2mL locking tubes (Fisher Scientific, 540295) with a ceramic bead on dry ice, powdered in a CryoMill (Retsch) cooled by liquid nitrogen, and weighed to 10-20mg into a new dry-ice-cooled 2mL locking tube and replaced on dry ice. Weight was recorded for later normalization.

Then, 700µL of ice cold 1mM sodium hydroxide was added to samples and vortexed for 30 seconds. 50µL of CoA internal standard mix consisting of a mixture of standard isotopically-labeled acyl-CoAs (see description in Standards and Chemicals above) was added to each calibration standard, quality control standard, and sample. 250µL of 250 mM sodium hydroxide was added to reach pH 12-13, and samples were vortexed. Samples were incubated in a 55°C heat block for two hours. 250µL of 1M Trizma-HCl was added to bring pH to 7 and samples were vortexed. 30µL of 100mM N-ethyl-maleimide was added to each sample to derivatize CoA; samples were vortexed and incubated overnight at room temperature. 50µL glacial acetic acid was added to bring pH to <4, and samples were vortexed. Samples were spun for 15 minutes at 21,300xG to remove proteins.

Supernatants were then purified via solid phase extraction using a vacuum manifold (2100-24, Analytical Sales and Services). A 2-(2-pyridyl)ethyl column (11-100-8971, Millipore Sigma) was equilibrated with 1 mL of 50% methanol with 2% acetic acid. Supernatant was applied, and column was washed twice with 1 mL of 50% methanol with 2% acetic acid and once with 1 mL of water. Samples were eluted into 10 mL glass centrifuge tubes (0553841C, DWK Life Sciences) with 2mL of 95% ethanol with 50 mM ammonium formate followed by 1mL of 90% methanol with 15 mM ammonium hydroxide. Samples were evaporated to dryness under a steady stream of nitrogen using a nitrogen evaporator manifold (11634, Organomation). Samples were resuspended in 100µL of water, centrifuged at 21,300xG for 5 minutes to remove insoluble particles, and transferred to MS vials (vials, Thermo Scientific, 6ERV11-03PPC; caps, Thermo Scientific, 6PRC11ST1R) for measurement.

### Mouse serum extraction for pantothenate LC-MS measurement

For serum samples analyzed in these studies, a methanol protein precipitation was performed as follows. Five µL of serum was transferred on wet ice to a 1.5 mL polypropylene microcentrifuge tube (Thermo Fisher Scientific, 3499). and 50 volumes (250 µL) of 100% MeOH cooled to -20°C was added. Samples were vortexed and incubated at −80°C overnight. Following incubation, samples were centrifuged at 21,300xG for 25 min at 4°C. Supernatants were transferred to MS vials (Thermo Scientific, 6ERV11-03PPC; caps, Thermo Scientific, 6PRC11ST1R) for measurement.

### LC-MS data acquisition

Mass spectrometry data was acquired on a high-resolution Q-Exactive Plus Orbitrap mass spectrometer equipped with a Vanquish Horizon UHPLC binary system and autosampler (Thermo Scientific). For total CoA analyses and untargeted high-resolution mass spectrometry analyses, metabolite extracts (5µL) were separated on a XBridge BEH Amide, 2.5 um, 2.1mm x 150 mm HILIC column (186006724, Waters) with XBridge BEH Amide XP VanGuard guard column (Waters 186007763) held at 25°C. LC separation was performed at 150µL/min using solvent A (95%:5% water:acetonitrile with 20 mM ammonium acetate, 20 mM ammonium hydroxide, 5uM ammonium phosphate, 0.5 µM methylenediphosphonic acid-acid, pH 9.45) and solvent B (100% acetonitrile) with the following gradient: 0 min, 90% B; 2 min, 90% B; 3 min, 75% B; 7 min, 75% B; 8 min, 70% B, 9 min, 70% B; 10 min, 50% B; 12 min, 50% B; 13 min, 25% B; 14 min, 25% B; 16 min, 0% B, 20.5 min, 0% B; 21 min, 90% B; 25 min, 90% B. The total gradient length was 26 minutes.

For total CoA experiments, LC-MS data was acquired in selected ion monitoring mode with a mass range of 850-950 m/z using electrospray ionization. The MS scans were in positive-ion mode with a resolution of 70,000 at m/z 200. The automatic gain control target was 3e6, and the maximum injection time was 200ms. Source ionization parameters were optimized with the spray voltage at 4.0 kV and in-source CID of 5.0eV, and other parameters were as follows: capillary temp at 425, S-lens level at 50, aux gas heater temperature at 400, sheath gas at 35, aux gas at 10, and sweep gas flow at 2.

For measurement of pantothenate in serum (used for normalization of total CoA labeling in Figure 4E and 4F), full MS analyses were acquired over a mass range of m/z 70-1000 using electrospray ionization and a method switching between positive and negative modes with a resolution of 70,000. The automatic gain control target was set at 3e6 ions, and the maximum injection time was 240 milliseconds. Source ionization parameters were optimized with the spray voltage at 3.0 kV, and other parameters were as follows: capillary temperature at 250, S-lens level at 50, heater temperature at 100, sheath gas at 40, aux gas at 10, and sweep gas flow at 0.

### Data analysis

Data were acquired using XCalibur (v.4.5.474.0; Thermo Scientific). To convert .raw files to mzML, we used ProteoWizard (https://proteowizard.sourceforge.io, version 3). Samples were analyzed using Skyline 4.1 (Michael MacCoss, University of Washington) and EL-Maven (v-0.12.1-beta), then data was analyzed and plotted with RStudio (2024.04.2+764), and GraphPad Prism (10.4.2).^28,29^ The ions at 893.1702 m/z and 897.1646 m/z representing unlabeled and labeled CoA-NEM were monitored with 15 ppm mass tolerance. Graphs were made using GraphPad Prism (10.4.2), and schematics were made using Biorender.

### Method Validation

The targeted HILIC–MS total CoA assay was evaluated according to guidelines established by the CLSI LC–MS C62-A document and the FDA bioanalytical method validation guidance for industry.^25,30^ Method parameters including the limit of quantification, linearity, accuracy, precision, carryover, stability, and matrix effect were evaluated, and results are shown in figures and associated text. All validation experiments were performed using acetyl-CoA standard (MedChemExpress, HY-113596A) in 60% fetal bovine serum pseudo-matrix, except for the matrix effect assessment, which was performed in 60% fetal bovine serum or mouse spleen tissue to compare extraction and ionization efficiencies between different matrices and validate use of the pseudo-matrix. For all studies, reagents were thawed and samples were prepared immediately prior to beginning hydrolysis/derivatization procedure. Samples were analyzed immediately after preparation, except for studies measuring autosampler stability in which samples were placed at 4°C for four days before analyzing.

Stocks of CoA standards were prepared by dissolving powder in 100% water (or DMSO for palmitoyl-CoA), and concentration was confirmed by measuring absorbance on a Beckmann DU640 spectrophotomer in fixed wavelength mode and blanking with pure solvent. The previously reported molar absorptivity of Coenzyme A at 259 nm, originating from the adenine ring, is independent of acyl chain and thus allowed us to calculate concentrations for the acyl-CoA standards (see absorbance values in Supplemental Table 1).^31^ Based on these concentrations, a working stock of 100µM was made. Calibration curves were prepared from the 100µM working stock of acetyl-CoA in 60% fetal bovine serum, diluted to 5µM, 25µM, 50µM, 75µM, and 100µM (which corresponded to 500-10,000 total picomoles per 100µL sample). A zero calibrator with no acetyl-CoA and only CoA internal standard mix was also measured. Peak areas for CoA-N-ethyl-maleimide (CoA-NEM) derivative (893.17017m/z) and CoA-NEM internal standard (897.16462 m/z) were calculated, and the ratio of CoA-NEM to CoA-NEM internal standard peak area was plotted against concentration. Curves were fit with a linear regression with 1/x weighting, and R^2^ of line of best fit and accuracy of calibrators is reported. For accuracy and precision studies, five independent samples of acetyl-CoA standard were made per quality control level (500 picomoles, 3000 picomoles, 6000 picomoles per 100µL sample). These 15 samples were prepared and run on three independent days, along with a freshly prepared calibration curve. The percent accuracy for each calibrator was calculated by comparing the true concentration to the value calculated from the calibration curve. For precision, the percent relative standard deviation of the five samples at a given calibrator level were calculated. For autosampler stability, these same samples at 500, 3000 and 6000 picomoles (n=5 each) were run immediately after preparation and after 4 days in the autosampler at 4°C.

For matrix effect studies, solutions of 100µL of 50µM acetyl-CoA were prepared in neat solution (water), 60% fetal bovine serum, and tissue matrix (mouse spleen, which has low endogenous CoA). Samples were prepared as described above and ratio of CoA-NEM to CoA-NEM internal standard was compared across conditions. For tissue precision study, five independent aliquots of tissue powder were weighed from the same tissue sample (mouse liver, brain, heart, or kidney). These were each hydrolyzed, reacted with NEM, and measured by LC-MS, and percent relative standard deviation was calculated as (standard deviation/mean) x 100.

### Total CoA Quantification in mouse tissues

The analytical workflow was applied to measure total CoA levels in tissues from 11-week-old male and female C57BL/6 mice. Tissue samples were prepared in batches of maximum 20 samples at a time, according to the protocol described above. To calculate tissue concentrations, all batches of samples were prepared and run with a fresh calibration curve, injected at the beginning and end of each instrument run. After comparing CoA-NEM/CoA-NEM internal standard ratio to calibration curve to establish the absolute number of picomoles in the sample, this value was normalized to the initial tissue powder weight in milligrams to give a picomole/milligram value for each sample.

### Analyzing total CoA synthesis in mouse liver

Mice were intravenously infused with [^13^C_3_-^15^N_1_] pantothenate as described above (Mouse Studies), and total CoA from livers was prepared and measured, without the use of [^13^C_3_-^15^N_1_] acyl-coA internal standard. Isotopic peaks were extracted, and AccuCor2 was used to account for natural isotopic abundance to calculate the percent of [^13^C_3_-^15^N_1_] pantothenate in serum and [^13^C_3_-^15^N_1_] CoA-NEM from [^13^C_3_-^15^N_1_] pantothenate in liver.^32^ Finally, liver CoA-NEM labeling was normalized to serum pantothenate labeling to calculate the newly-synthesized fraction of total CoA.

## Results & Discussion

### Sample preparation and derivatization strategy for total CoA measurement

Measuring Coenzyme A concentration and synthesis rate from tissues is scientifically valuable yet technically challenging. Coenzyme A acts as a cofactor in several metabolic pathways, including the tricarboxylic acid cycle, fat synthesis, and fat oxidation (Figure 1A). The concentration of Coenzyme A in cells governs the ability to carry out these pathways. However, acyl-Coenzyme A species are challenging to measure: acyl chains of different lengths impart different biochemical properties, making it challenging or impossible to capture all acyl-CoA species with the same analytical method, and some acyl-CoA species are labile or present at low concentrations. In addition, measuring CoA synthesis rate using isotope tracing is only possible using mass spectrometry, while many previous CoA-measurement methods use other approaches such as HPLC-UV/Vis spectroscopy.^12–24^

Therefore, we set out to develop a liquid chromatography-mass spectrometry (LC-MS) method to measure total Coenzyme A, defined as the sum of CoASH plus acyl-CoA concentrations. To measure total CoA concentration in tissues, we ground flash-frozen mouse tissues to powder using a liquid-nitrogen-cooled CryoMill, added [^13^C_3_-^15^N_1_]-labeled yeast-derived acyl-CoA internal standard to correct for process variability, used base and heat to hydrolyze acyl chains, and derivatized the resulting CoASH with N-ethyl-maleimide (Figure 1B, see detailed description in Methods). The Michael addition with N-ethyl-maleimide (NEM) protected the thiol of CoASH from conjugation with other cellular thiols like glutathione.^33–35^ Then, after solid phase extraction and reconstitution, we performed liquid chromatography-mass spectrometry using hydrophilic interaction chromatography (Figure 1C).

### Efficiency of basic hydrolysis and N-ethyl-maleimide derivatization of acyl-CoAs

We validated this hydrolysis and derivatization method first using pure acetyl-, succinyl-, and palmitoyl-CoA standards to verify conversion of acyl-CoAs to CoA-NEM. The method displayed excellent hydrolysis and derivatization efficiency for acetyl-CoA and succinyl-CoA standards, with 1000-fold reduction in acyl-CoA levels and corresponding increase in CoA-NEM (chromatograms in Figure 2A, raw peak areas in Supplemental Figure S1). Note that succinyl-CoA standard was measured as a combination of succinyl-CoA and CoASH, likely due to its marked instability in solution (half-life reported as 1-2 hours).^36,37^ Hydrolysis and derivatization of a palmitoyl-CoA standard yielded a similar CoA-NEM peak intensity (Supplemental Figure S2), although we are unable to detect the intact species prior to hydrolysis with the LC-MS method used here. Therefore, this total CoA method captures the CoA moiety from both short and long chain acyl-CoA species. We also verified that our [^13^C_3_-^15^N_1_]-labeled acyl-CoA internal standards underwent NEM derivatization as expected, resulting in a [^13^C_3_-^15^N_1_]-CoA-NEM peak that co-elutes with the unlabeled CoA-NEM analyte (Supplemental Figure S3).

**Figure 2.**
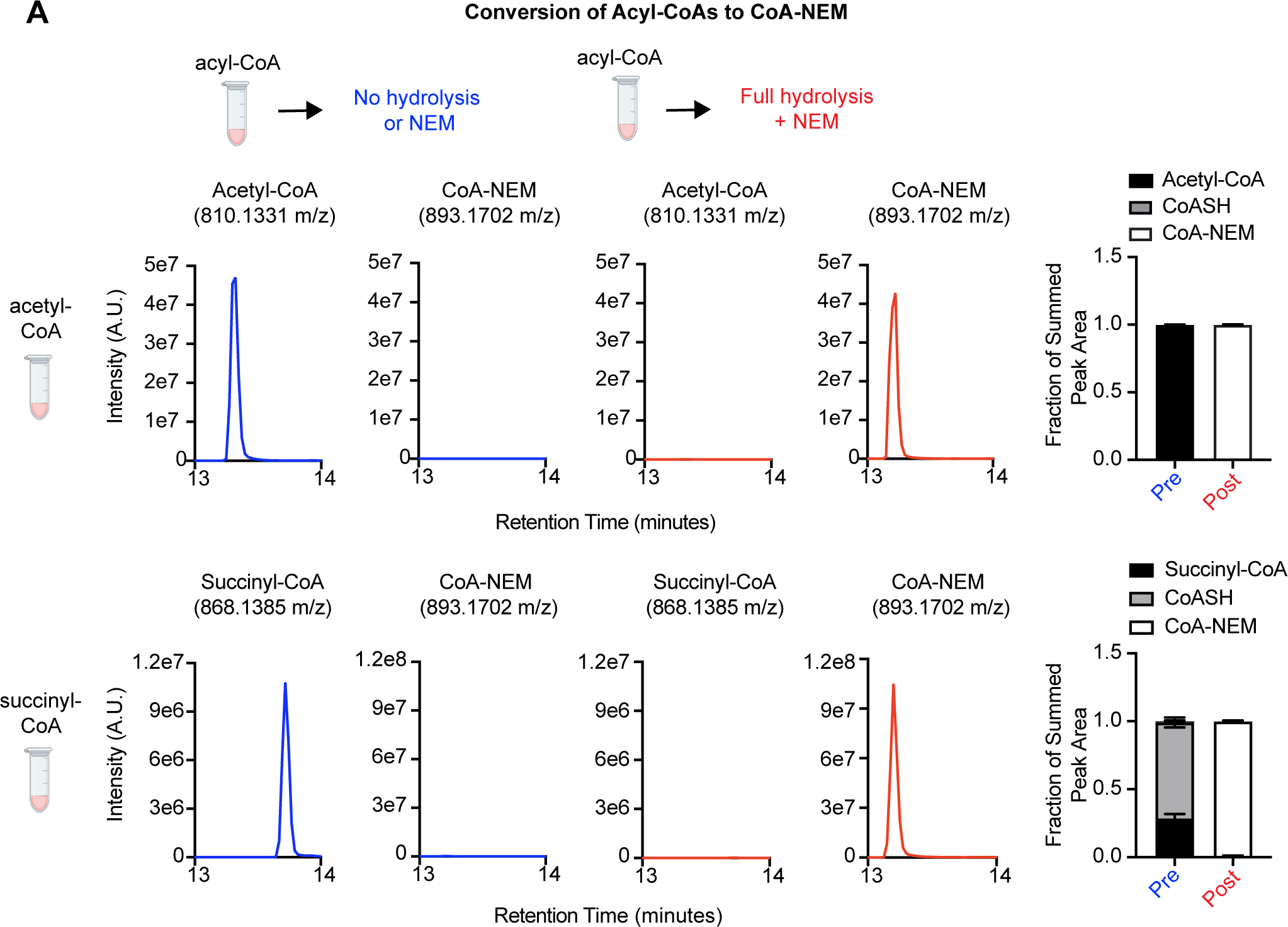
Efficiency of basic hydrolysis and N-ethyl-maleimide derivatization of acyl-CoAs. **A)** Representative chromatograms of acyl-CoAs (acetyl-CoA, top and succinyl-CoA, bottom) and CoA-NEM before (blue, left) and after (red, center) hydrolysis/derivatization procedure using 50 µM neat standards. Bar graphs (far right) show peak areas of chromatograms on the left for CoA species detected pre- (blue) and post- (red) hydrolysis/derivatization procedure from n=2 50µm neat standards, displayed as mean with standard deviation.

### Validation and key figures of merit for total Coenzyme A measurement method

To validate our total CoA measurement method, we evaluated its linearity, accuracy and precision. Calibration standards consisted of 100 µL solutions of acetyl-CoA prepared at 5, 25, 50, 75, and 100 µM, representing 500-10,000 picomoles per sample, with 25 to 500 picomoles on column after a 5 µL injection. Utilizing 5 µL injections of these calibration standards, the peak areas of CoA-NEM and CoA-NEM internal standard were calculated for each calibrator, and the ratio of CoA-NEM to CoA-NEM internal standard peak area was plotted against concentration. Utilizing a linear regression with 1/x weighting, the method displayed excellent linearity, with all calibrators falling well beneath 15% error as recommended by the FDA’s Bioanalytical Method Validation Guidelines, and displaying R^2^ > 0.99 in three independent runs (Figure 3A-B).^25^ This calibration and weighting strategy was validated with independently prepared quality control samples made from separate acetyl-CoA stocks at three concentrations spanning the linear range (500 picomoles, 3000 picomoles, and 6000 picomoles per sample), which displayed excellent accuracy (calculated as percent error) and precision (calculated as relative standard deviation) in three independent runs prepared and run on three separate days (Figure 3C). Additionally, samples were stable over 96 hours in a 4°C autosampler (Figure 3D), and carryover from the highest concentration calibrator was <5% signal of the lowest calibrator (Supplementary Figure S4).

**Figure 3.**
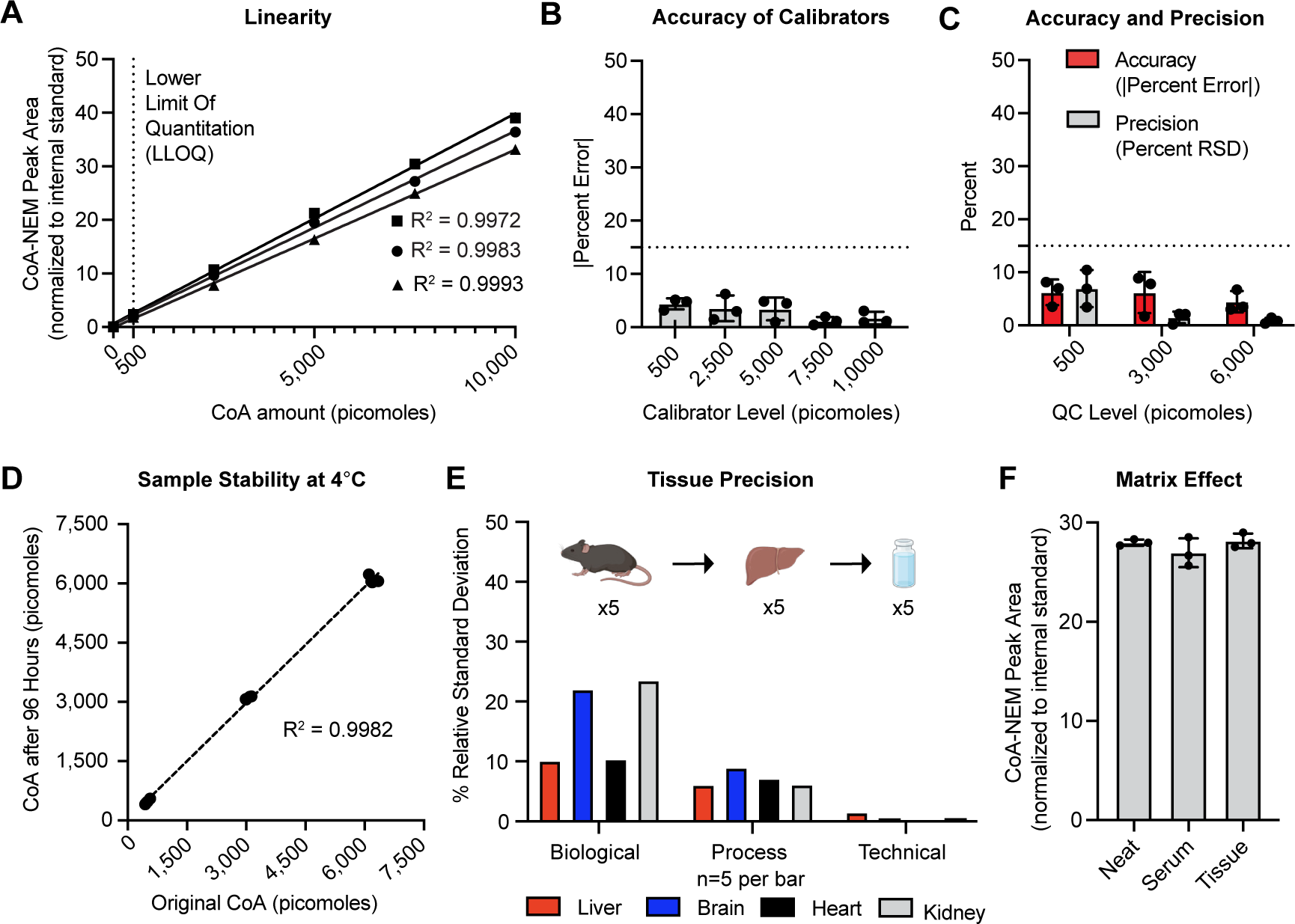
Validation and key figures of merit for total Coenzyme A measurement method. **A)** Linearity studies from n=3 independent runs (represented by different shapes) with 5 non-zero calibrators and a zero calibrator prepared from acetyl-CoA standards in 60% fetal bovine serum pseudo-matrix. Data was fitted with linear regression with 1/x weighting, and R^2^ value is shown for each independent run. **B)** Accuracy of calibrators shown in **A)**, prepared by calculating the percent error from the line of best fit to the known concentration. Data is shown as mean and standard deviation from n=3 independent runs. Dotted line indicates 15%, the maximum permissible error for method validation. **C)** Summary data from n=3 independent accuracy and precision studies with n=5 independently prepared acetyl-CoA standards samples at each quality control level in each run. Each accuracy point represents the absolute value of the average percent error for the 5 calibrators within a run. Each precision point represents the %RSD between the 5 calibrators at a given concentration within a given run. Data shown as mean and standard deviation. **D)** Autosampler stability showing calculated concentration of n=5 acetyl-CoA standards at 500, 3000 and 6000 picomoles run immediately after preparation and after 4 days in autosampler held at 4°C. Linear regression between original and rerun samples shows R^2^ >0.99. **E)** Tissue precision (shown as percent RSD) from n=5 biological replicates, n=5 process replicates (separately prepared samples of powder from the same tissue sample), and n=5 technical replicates (same sample vial injected five times on LC-MS). **F)** n=3 process replicates of 50µM standard (5,000 picomoles) prepared neat, in 60% serum matrix, or in spleen tissue matrix. Data shown as mean and standard deviation.

We further validated the precision of our method on multiple mouse tissue types, as extraction from tissue can introduce metabolite loss and inter-sample variation. For these studies, we analyzed either n=5 of the same tissue type collected from different mice to assess biological variability, n=5 aliquots of frozen tissue powder from the same mouse (process variability), or n=5 injections of the same processed sample in the LC-MS (technical variability). We found that our method displayed excellent precision (<15% relative standard deviation) for repeated preparations of the same tissue sample (process variability, Figure 3E), highlighting the method’s reproducibility in extracting CoA from tissue. As expected, biological variability was greater than process variability, since it encompasses both differences between individual mice plus the process variability produced during sample preparation.

Note that because Coenzyme A is present in all cells, calibrators and quality control samples could not be prepared in the same matrix (mouse tissue) as unknown samples. Because the levels of CoA are much lower in blood serum than tissue matrices, fetal bovine serum was chosen as a pseudo-matrix due to ease of access and minimal detectable endogenous CoA pool (Supplemental Figure S5).^38^ The amount of fetal bovine serum used (60% in water) was selected to match the protein concentration of tissue samples, measured by BCA protein assay (Supplemental Figure S6). To evaluate matrix effect, the same concentration of acetyl-CoA standard was either prepared neat in water, prepared in 60% fetal bovine serum, or spiked into mouse spleen homogenate, a tissue with relatively low CoA level. No significant matrix effect was observed (<5% alteration in peak area ratio and no retention time shift) across neat standard or standard in serum or tissue matrix, validating the use of serum as a pseudo-matrix (Figure 3F).

### Comparison to previous CoA measurement methods

Our CoA hydrolysis-derivatization approach can quantify both total CoA concentration and isotopic labeling, which previous methods could not. The earliest methods to quantitate acyl-CoA species took advantage of specific enzymatic reactions utilizing CoASH or acetyl-CoA that generate products measurable by spectroscopy or radioactivity detection. For example, enzymatic-spectroscopic methods react malonate decarboxylase with acetyl-CoA followed by addition of acetate kinase and absorbance of resulting acetohydroxamate, which allows for measures of acetyl-CoA, malonyl-CoA, and CoASH (via various modifications of the procedure).^12,13,15^ Similarly, radioassays reacted acetyl-CoA with [^14^C]-oxaloacetate or CoASH with [^14^C]-palmitic acid, products of which are then measured by scintillation counter.^16,18,39^ These methods, while sensitive, lack the ability to capture the total CoA content of the cell, instead reporting a summed measurement of the most abundant short-chain CoA species. Additionally, these methods are unable to distinguish stable-isotope-labeled CoA from unlabeled CoA.

More recent methods use high-performance liquid chromatography-ultraviolet detection (HPLC-UV) or HPLC-fluorescence detection to measure total CoA concentration. These methods hydrolyze acyl-CoAs before reacting CoASH with a UV- or fluorescence-active probe.^19–24,40^ These methods are similar to the method described herein in their ability to capture total CoA pools, but they cannot measure CoA isotopic labeling. Thus, they cannot measure CoA synthesis rate using stable-isotope tracing and cannot use heavy internal standards to assist in absolute CoA quantification. Additionally, there are HPLC-UV and LC-MS/MS methods that measure specific acyl-CoA species and their isotopic labeling, but due to the large differences in chemical properties between short and long chain CoA species, none of these methods are able to fully capture the total CoA pool with one separation method. ^38,41^

### Quantification of total Coenzyme A concentration and synthesis rate in mouse tissues

Using the method reported, we quantified total CoA levels in liver, heart, brain, kidney, and pancreas of male and female mice (Figure 4A). Values ranged from around 30 pmol/mg in pancreas to around 550 pmol/mg in liver, spanning a range of over 10-fold as has been reported previously.^42^ Note that since 10 to 20 milligrams was used for each tissue sample measurement, these samples had roughly 500 to 10,000 picomoles of CoA per 100µl sample after preparation, falling in the linear range for our method (Figure 4A, Figure 3A). Consistent with prior literature, liver and heart have the highest total CoA concentration across tissues.^42^ Figure 4B compares the total CoA concentrations measured here to previous published CoA measurements (also see Table 1). Many past studies measured only heart or liver CoA, and so far we have not found any measuring mouse pancreas.^12–14,16,18–24,38–41^ Our measured total CoA concentrations are higher than prior measurements: for example, our liver total CoA concentration is 2-4-fold higher than previously published values. Possible contributing factors may include our rigorous tissue preparation including freeze-clamping and maintaining tissues at dry-ice or liquid-nitrogen temperature until extraction; the fact that we directly capture the total CoA pool, while most previous methods instead sum concentrations for only a few short-chain acyl-CoA species; and our ability to account for CoA loss during sample preparation with use of internal standards. However, note that published CoA concentrations vary widely within and between methods, making it challenging to directly compare results obtained from separate studies.

**Figure 4.**
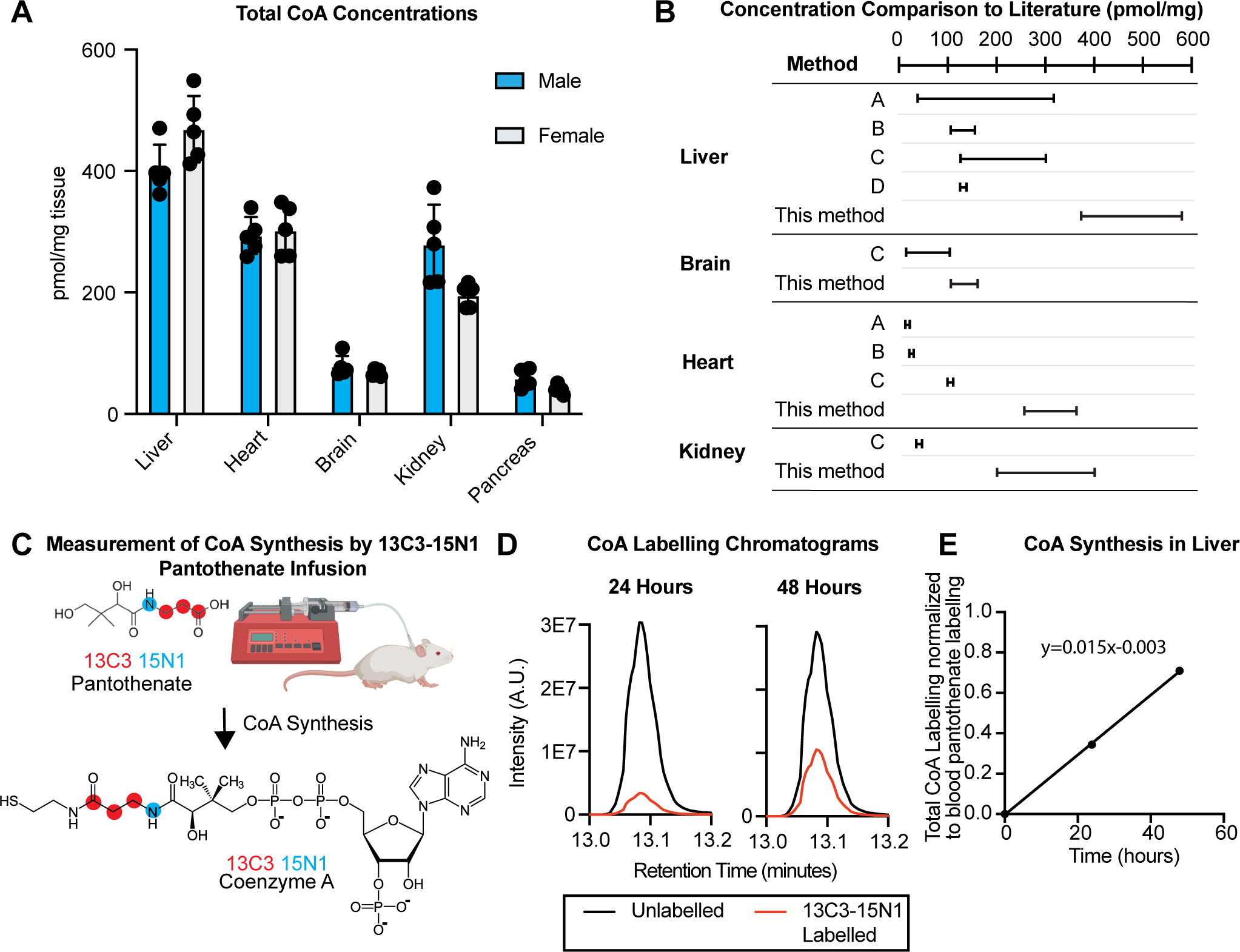
Quantification of total Coenzyme A concentration and synthesis rate in mouse tissues. **A)** Total CoA levels in tissues of n=5 male and female mice after a 9-hour daytime fast. Data shown as mean and standard deviation. **B)** Comparison of literature values for total CoA concentrations in tissue. Concentrations in pmol/mg for given tissues of interest, shown as ranges reported for a particular method class (A: enzymatic spectroscopy, B: enzymatic radioassay, C: HPLC-UV/Vis or HPLC-FLD, D: LC-MS/MS) compared to the method detailed herein. Note, classes A, B, and D’s values are a summed measurement of the reported species, rather than a true total CoA measurement. **C)** Infusion schematic for stable isotope tracing of pantothenate into Coenzyme A to measure of CoA synthesis rate in live awake mice. **D)** Representative chromatograms showing labeled and unlabeled CoA-NEM from heavy-labeled pantothenate infusion studies. Intensity shown in A.U. (arbitrary units). **E)** Ratio of labeled CoA normalized to blood serum pantothenate labeling in the same mouse, with a linear fit representing the rate of synthesis of new CoA from pantothenate. N=1 mouse per timepoint.

**Table 1.**
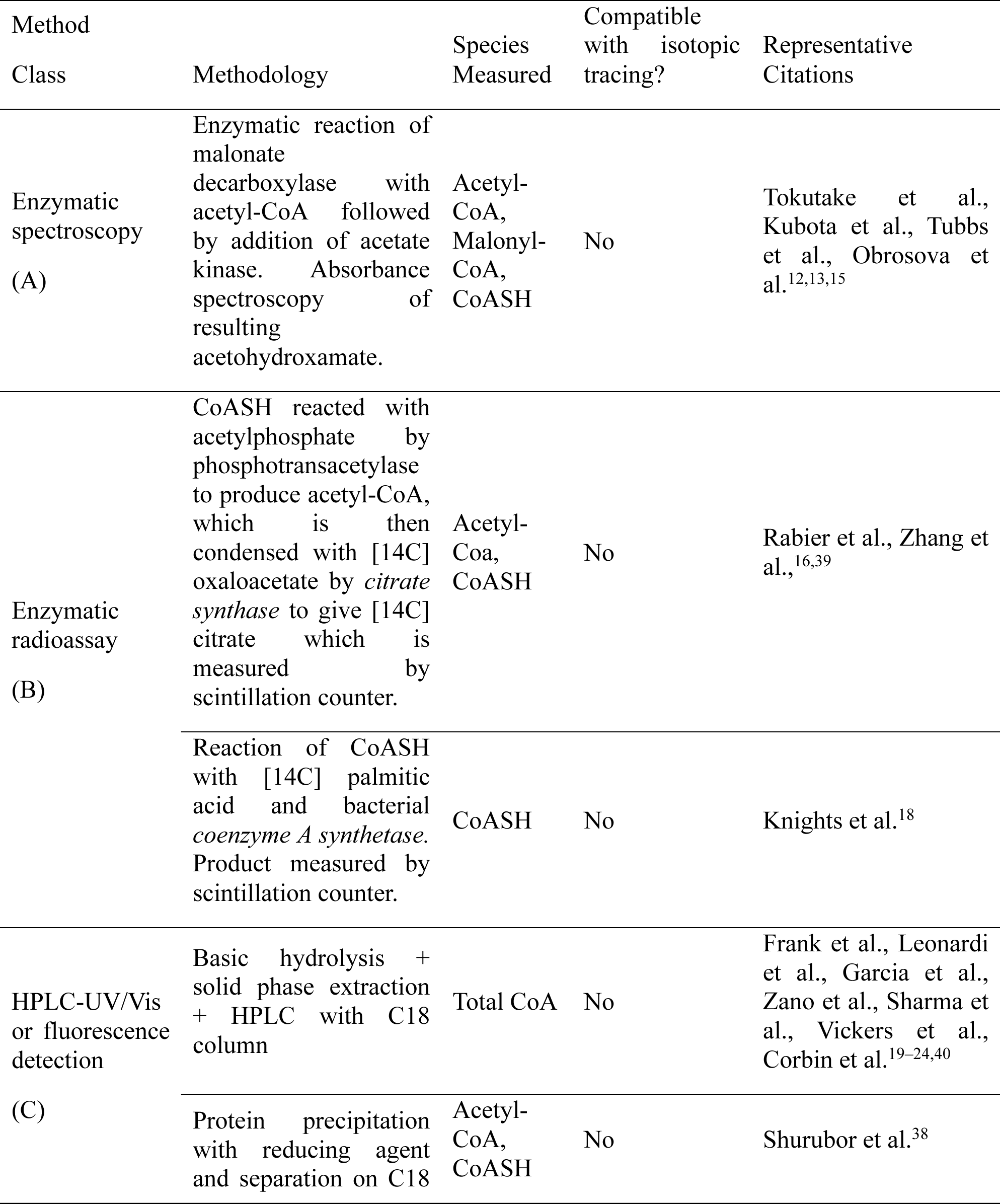
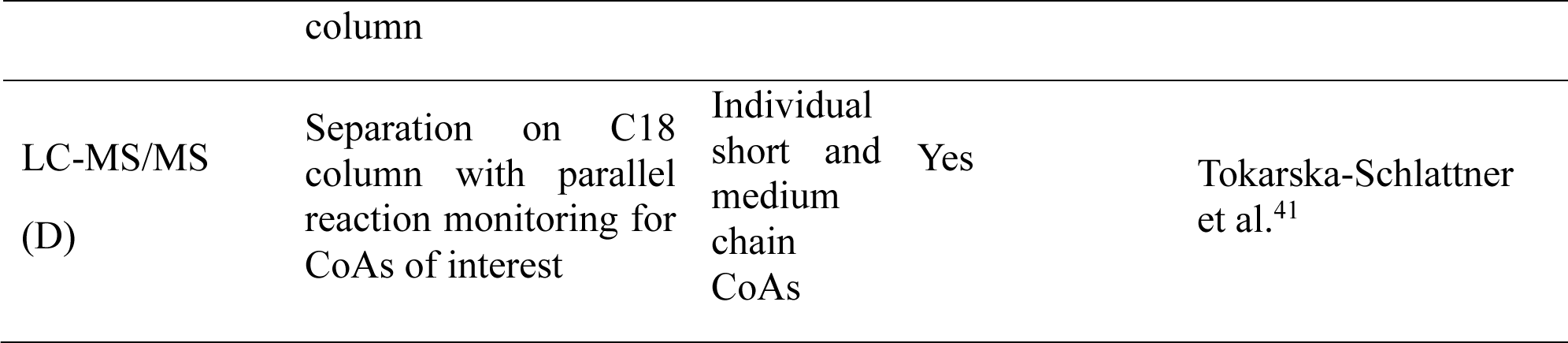
Historical methodologies used for quantification of total Coenzyme A or individual acyl-Coenzyme A species.

This LC-MS total CoA measurement method, unlike enzymatic or fluorescent methods used previously, can distinguish isotope-labeled CoA from endogenous CoA. This enables use of isotope-labeled acyl-CoA internal standards as described above. Further, it enables use of stable isotope tracing using CoA precursors like [^13^C_3_-^15^N_1_] pantothenate to measure CoA synthesis (note that in this application we omit the heavy internal standards from the workflow as the internal standards cannot be distinguished from the tissue CoA made from heavy pantothenate). To measure CoA synthesis in mouse tissues, we intravenously infused [^13^C_3_-^15^N_1_] pantothenate in unrestrained, unanesthetized mice for 24 or 48 hours (Figure 4C). We used our total-CoA method to measure the enrichment of [^13^C_3_-^15^N_1_] CoA-NEM in mouse livers, and normalized to the enrichment of the precursor [^13^C_3_-^15^N_1_] pantothenate in blood serum to calculate the fraction of CoA synthesized from blood pantothenate (Supplemental Figure S7).^43^ We found that the fraction of mouse liver CoA newly synthesized from pantothenate reaches 35% at 24 hours and 70% at 48 hours (Figure 4D-E). When fitting a linear regression to the plot of labeled fraction against time, the labeling increases at a rate of 1.5% per hour, so that 50% of tissue CoA is turned over in about 33 hours. This is consistent with an *in vitro* measurement of CoA synthesis where roughly 10% of CoA turned over in 5 hours (i.e. 2% per hour).^2^

### Conclusions

Here, we developed a method to quantify total Coenzyme A concentration and isotopic labeling using LC-MS. This method is accurate and precise. Further, it offers the ability to measure isotopic forms of CoA, unlocking the ability to use heavy internal standards and to measure total CoA synthesis rate. We applied this method to quantify total CoA concentrations in five tissue types in male and female mice, and to quantify CoA synthesis rate in mouse liver.

## Supporting information

All Supplemental Figures

## ASSOCIATED CONTENT

### Supporting Information

The following files are available free of charge.

Supplemental Figures (PDF): Concentration of CoA standard solutions, peak areas for pre- and post- hydrolysis and derivatization samples, hydrolysis and derivatization of long-chain CoA and internal standards, carryover results, blank matrix protein concentration and CoA signal, labeling from infusion study

## AUTHOR INFORMATION

### Author Contributions

ALT, CRB, and NWS conceptualized the study and contributed to data interpretation. ALT performed the experiments and data analysis for the study. CRB supervised the project. The manuscript was written through contributions of all authors. All authors have given approval to the final version of the manuscript.

### Funding Sources

This work was supported by R00CA273517 and V Foundation V2024-009 to CRB, T32GM071339 to ALT, and R35GM156596 to NWS.

## ACKNOWLEDGMENTS

The authors thank Trevor Penning and Catia Marques for use of the UV/Vis spectrometer.

## ABBREVIATIONS

CoA: Coenzyme A;
NEM: N-ethyl-maleimide;
FLD: fluorescence detection;
HPLC-MS: high performance liquid chromatography-mass spectrometry;
UV/Vis: ultraviolet/visible

## Notes

### Competing Interest Statement

The authors have declared no competing interest.

