## Supplementary material for "A liquid chromatography-mass spectrometry method to quantify total Coenzyme A concentration and isotopic labeling": All Supplemental Figures

**Table of Contents**

**P2**—Table S1: Acyl-CoA standard concentrations

**P3**—Figure S1: Pre and post hydrolysis and derivatization peak areas for acyl-CoA and CoA-NEM

**P4**—Figure S2: Hydrolysis and derivatization of short and long chain acyl-CoA

**P5**—Figure S3: Hydrolysis and derivatization of stable isotope labeled acyl-CoA

**P6**—Figure S4: Carryover

**P7**—Figure S5: Blank pseudo-matrix CoA signal

**P8**—Figure S6: Protein concentration in pseudo-matrix versus tissue matrix

**P9**—Figure S7: Raw labeling in stable isotopically-labeled pantothenate infusion study

**Supplemental Table S1. Absorbance of acyl-CoA standard solutions by UV-Vis absorption spectrophotometry and resulting calculated concentrations.**

| Standard | Vendor and Catalog Number | Absorbance of 1:200 diluted stock at 259nm | Calculated Stock Concentration |
|---|---|---|---|
| Palmitoyl-CoA | Avanti Polar Lipids 870716P | 0.2735 | 3.8177 mM |
| Acetyl-CoA | MedChemExpress HY-113596A | 0.3869 | 5.4 mM |
| Succinyl-CoA | MedChemExpress HY-148285 | 0.2765 | 3.859 mM |

**Supplemental Figure S1. Integrated peak areas before and after hydrolysis and derivatization of acetyl- and succinyl-CoA, corresponding to Figure 2.** N=2 independent samples without (blue) and with (red) hydrolysis and NEM derivatization procedure on 50µM acetyl-CoA (left) and 50 µM succinyl-CoA (right) standards.

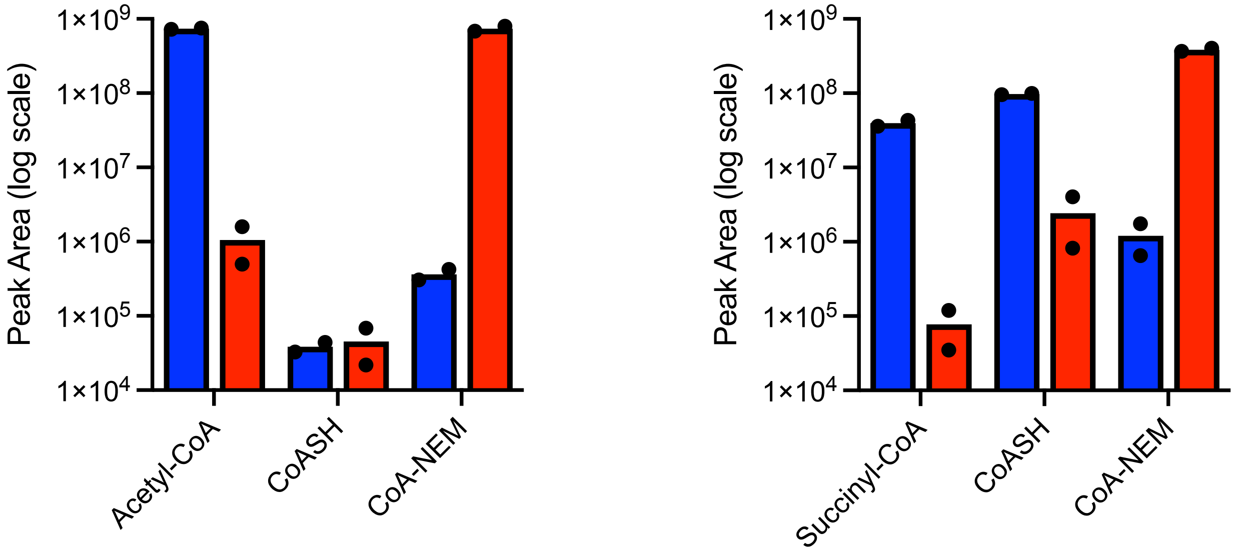

**Supplemental Figure S2. Comparative hydrolysis and derivatization between short- and long- chain CoAs. Left)** Representative chromatogram for CoA-NEM from hydrolysis and NEM derivatization of 50 µM palmitoyl-CoA standard. **Right)** Peak areas for CoA-NEM from hydrolysis and NEM derivatization procedure carried out on 50 µM standards of acetyl-CoA, succinyl-CoA, or palmitoyl-CoA (n=2 per condition). Data shown as mean and standard deviation.

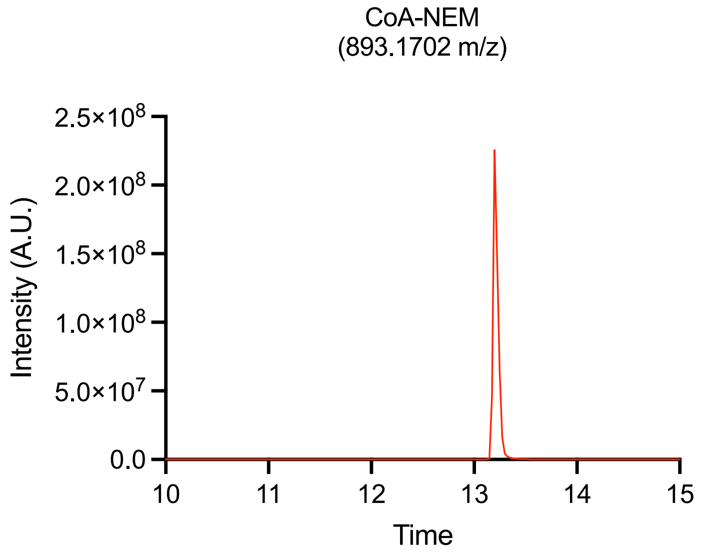


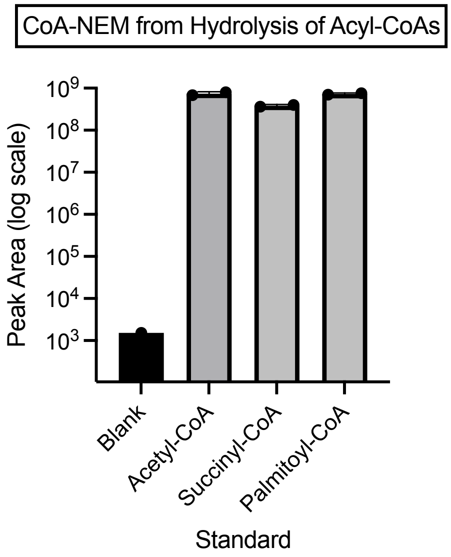

**Supplemental Figure S3. Hydrolysis and derivatization of stable isotopically labeled CoA internal standard mix.** Representative chromatograms for acetyl-CoA and CoA-NEM without (blue) and with (red) hydrolysis and NEM derivatization procedure, from 50µM acetyl-CoA standards with acyl-CoA heavy isotopically labeled internal standard. Top shows unlabeled standard, and reflection plot shows [$^{13}C_3$-$^{15}N_1$] labeled.

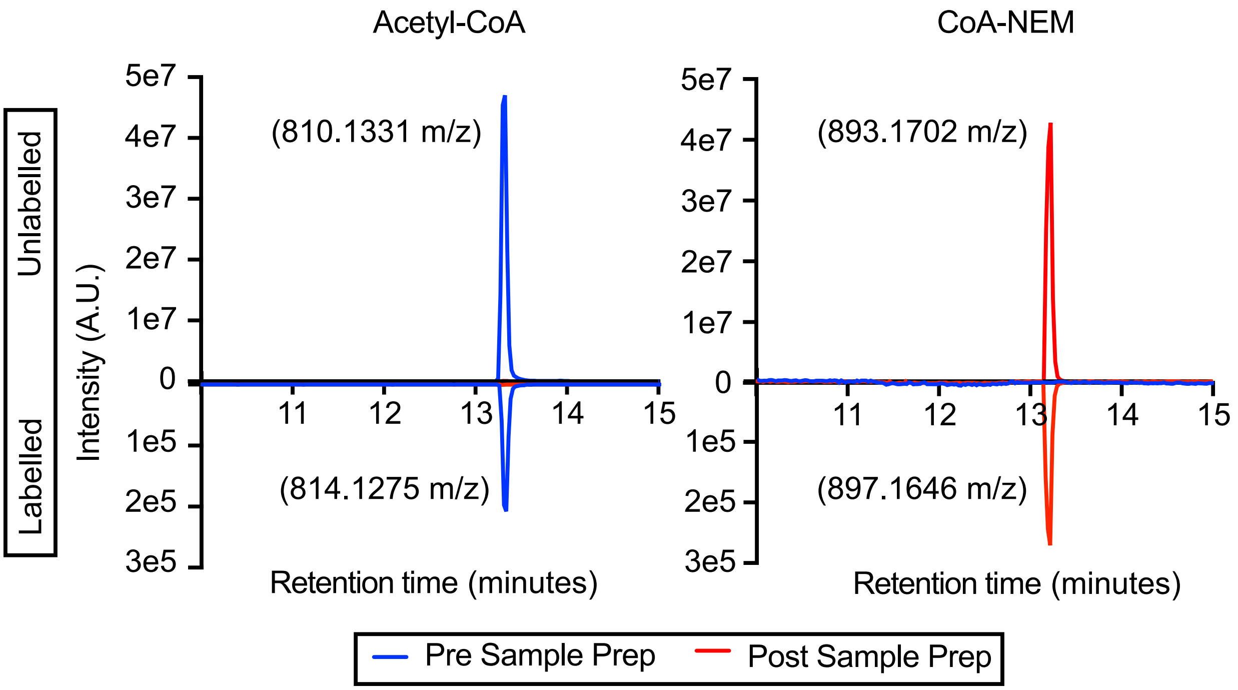

**Supplemental Figure S4. Validation of minimal carryover from analysis of hydrolyzed and derivatized acetyl-CoA.** Raw peak area of n=3 blanks (injected post highest calibrator) and lowest concentration calibrator. Data shown as mean and standard deviation, plotted on a log scale.

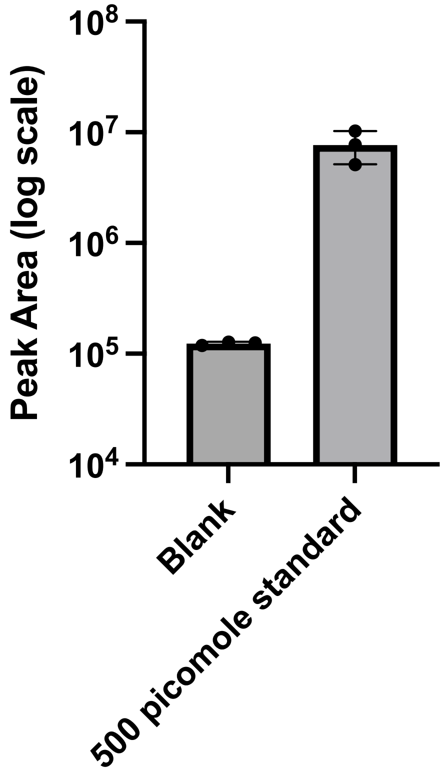

**Supplemental Figure S5. CoA-NEM signal from hydrolyzed and derivatized blank serum.** Raw peak area of 60% fetal bovine serum with no added CoA standard and lowest concentration calibrator. N=3 per bar, data shown as mean and standard deviation, plotted on a log scale.

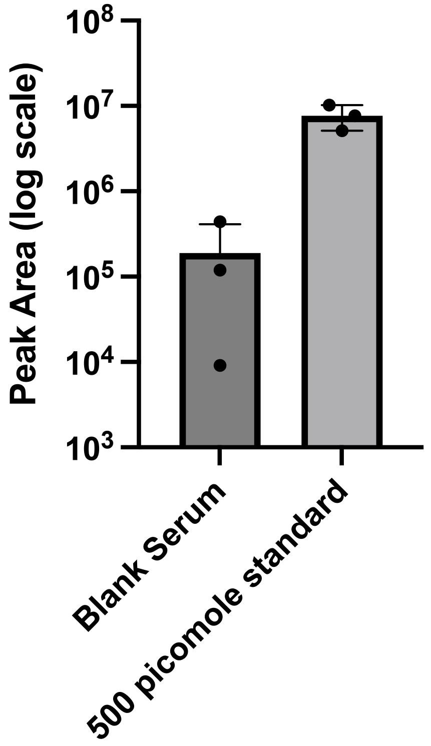

**Supplemental Figure S6. Protein concentration in true matrix (tissue) versus pseudo-matrix (serum) used to determine concentration of serum in calibrators and quality control samples.** Protein concentration by bicinchoninic acid assay for n=3 aliquots of fetal bovine serum and n=3 mouse liver tissue samples. Data shown as mean and standard deviation.

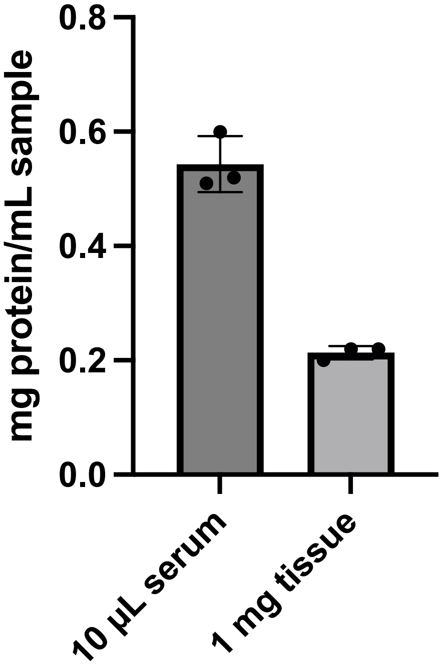

**Supplemental Figure S7. Labeling for serum pantothenate and for liver CoA-NEM from pantothenate intravenous infusion, used to calculate serum-normalized liver CoA-NEM labeling shown in Figure 4E.** [$^{13}C_3$-$^{15}N_1$] labeled fraction of total for serum pantothenate (black bar) and mouse liver tissue CoA-NEM (gray bar) from mice infused for 24 or 48 hours with [$^{13}C_3$-$^{15}N_1$] labeled pantothenate. n=1 per bar.

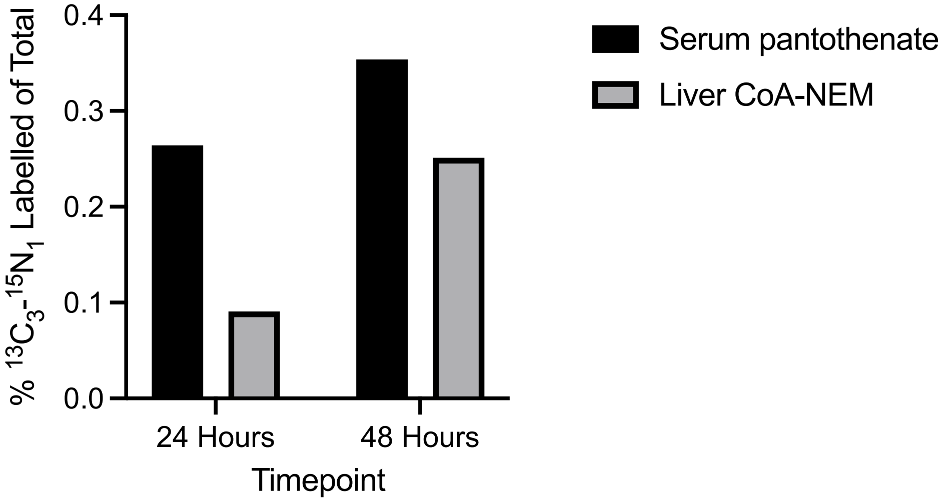
